# Eye-movement intervention enhances extinction via amygdala deactivation

**DOI:** 10.1101/282467

**Authors:** Lycia D. de Voogd, Jonathan W. Kanen, David A. Neville, Karin Roelofs, Guillén Fernández, Erno J. Hermans

**Author notes:** Correspondence to: Lycia D. de Voogd, Donders Institute for Brain, Cognition and Behaviour (Radboudumc), P.O. Box 9101, 6500 HB Nijmegen, The Netherlands, Fx.

## Abstract

Improving extinction learning is essential to optimize psychotherapy for persistent fear-related disorders. In two independent studies (both n=24), we found that goal-directed eye movements activate a dorsal fronto-parietal network and transiently deactivate the amygdala. Connectivity analyses revealed this down-regulation engages a ventromedial prefrontal pathway known to be involved in cognitive regulation of emotion. Critically, when eye movements followed memory reactivation during extinction learning, this reduced spontaneous fear recovery 24 hours later. Stronger amygdala deactivation furthermore predicted a stronger reduction in subsequent fear recovery after reinstatement. In conclusion, we show that extinction learning can be improved with a non-invasive eye-movement intervention that triggers a transient suppression of the amygdala. Our finding that another task which taxes working memory leads to a similar amygdala suppression furthermore indicates that this effect is likely not specific to eye movements, which is in line with a large body of behavioral studies. This study contributes to the understanding of a widely used treatment for traumatic symptoms by providing a parsimonious account for how working memory tasks and goal-directed eye movements can enhance extinction-based psychotherapy, namely through neural circuits similar to those that support cognitive control of emotion.

**Significant statement:** Fear-related disorders represent a significant burden on individual sufferers and society. There is a high need to optimize treatment, in particular via *non-invasive* means. One potentially effective intervention is execution of eye movements following trauma recall. However, a neurobiological understanding of how eye movements can reduce traumatic symptoms is lacking. We demonstrate that goal-directed eye-movements, like working memory tasks, deactivate the amygdala, the core neural substrate of fear learning. Effective connectivity analyses revealed amygdala deactivation engaged dorsolateral and ventromedial prefrontal pathways. When applied during safety learning, this deactivation predicts a reduction in later fear recovery. These findings provide a parsimonious and mechanistic account of how behavioral manipulations taxing working memory and suppress amygdala activity can alter retention of emotional memories.

## Introduction

Extinction learning is core to most effective therapies for disorders of fear and anxiety (Bisson et al., 2013). Exposure therapy, for instance, results in the formation of an extinction memory that suppresses fear expression. Relapse of pathological fear is nevertheless common (Maren, 2011; Dunsmoor et al., 2015b). Improving extinction learning is therefore an important goal of translational research into fear-related disorders (Dunsmoor et al., 2015b). Pharmacological treatments have proven effective in preventing fear recovery in animal models (Nader et al., 2000), but these methods are often not applicable in humans (Nader et al., 2000) or have yielded inconsistent results in experimental models with humans (Kindt et al., 2009; Bos et al., 2012). New non-invasive techniques have been developed that target reconsolidation of the original memory rather than enhance standard extinction learning (Schiller et al., 2010). Although these results are promising, their clinical utility is so far unclear.

Clinically effective treatments are not always derived from such experimental models. One example is Eye Movement Desensitization and Reprocessing (EMDR; Bisson et al., 2013), an evidence-based therapy and part of mental health care guidelines in many countries (Bisson et al., 2013; Lee and Cuijpers, 2013). Despite its wide use, a mechanistic, neurobiological understanding of EMDR is lacking. During treatment, patients divide their attention between recalling traumatic memories and making lateral eye movements directed by the therapist’s hand. Eye movements are central to the procedure, but it is unclear if they play *any* role in the therapeutic outcome above normal extinction (Lee and Cuijpers, 2013). Moreover, it is unclear whether the eye movements enhance extinction (i.e., new learning) or enhance the updating of the original memory trace (i.e., unlearning). Insight into the potential role of eye movements and the neurobiological mechanisms underlying this manipulation is not only crucial to further optimize this therapy but would also importantly advance our fundamental understanding of extinction learning.

One lead into a neural mechanistic account is that goal-directed eye movements are associated with activations in the dorsal fronto-parietal network, including frontal eye fields (FEF; Corbetta and Shulman, 2002), similar to working memory tasks that require goal-directed attention. Critically, working memory tasks are accompanied by robust deactivations in a posterior-medial network (Qin et al., 2009), including the amygdala. Interestingly, a similar activation of the dorsal fronto-parietal network and deactivation of the amygdala has been found during cognitive regulation of emotion as well (Ochsner and Gross, 2005). This is important because targeting the amygdala following memory reactivation, by blocking protein synthesis, prevents fear recovery in rodents (Nader et al., 2000). Similarly, systemic administration of propranolol, a *β*-adrenoceptor antagonist, presumably exerts its effects on fear recovery via inhibition of protein synthesis in the amygdala (Dȩbiec and Ledoux, 2004). Amygdala reactivity measured with BOLD-fMRI in humans is furthermore decreased after propranolol administration (Hurlemann et al., 2010). Indeed, it has been shown that working memory-like tasks, such as a game of Tetris (Holmes et al., 2009; James et al., 2015) can affect the emotionality of memories. We therefore hypothesized (1) that goal-directed eye movements could be used as a non*-*invasive tool to transiently suppress amygdala activity, comparable to working memory tasks, and (2) that a well-timed application of this deactivation following memory reactivation could reduce fear recovery.

To test our hypotheses, in experiment one (n = 24), participants performed a two-back working memory task and goal-directed eye movements in a block design while undergoing functional MRI. We tested whether both tasks would suppress amygdala activity as well as alter the coupling between the amygdala and dorsal lateral prefrontal cortex. In the second experiment (n = 24), we integrated eye movements into an established Pavlovian fear conditioning / extinction / recall paradigm and tested whether goal-directed eye movements prevent fear recovery via this amygdala deactivation.

## Materials and Methods

### EXPERIMENT 1

#### Participants

Twenty-four right-handed healthy volunteers (12 females, 12 males; 23-37 years [M=26.95, SD=3.6]) completed the study. Exclusion criteria were any contraindications for MRI. All gave written informed consent and were paid for their participation. This study was approved by the local ethical review board (CMO region Arnhem-Nijmegen).

#### Experimental design

The tasks consisted of 6 blocks of a two-back working memory task (Qin et al., 2009), 6 blocks of smooth-pursuit lateral eye movement task, and an additional 8 blocks of low-level fixation baseline. The duration of each block was 27 seconds. Within each two-back block, participants saw a random sequence consisting of 15 single digits. Each digit was presented for 400 ms, followed by an interstimulus interval (ISI) of 1400 ms. Participants were asked to detect whether the current item had appeared two positions back in the sequence and were instructed to make a button press when detecting a target. For the eye-movement blocks, participants were instructed to follow a laterally moving dot with their eyes. The speed of the eye movements was ∼1 Hz, based on previous laboratory models of EMDR (van den Hout et al., 2013). For 15 participants, eye-tracking data were recorded and verified participants complied with the instructions.

#### MRI data acquisition

MRI scans were acquired using a Siemens (Erlangen, Germany) MAGNETOM Skyra 3T MR scanner. T2*-weighted blood oxygenation level-dependent (BOLD) images were recorded using a customized EPI sequence with ascending slice acquisition (37 axial slices; TR, 1.89 s; TE, 25 ms; Generalized Autocalibrating Partially Parallel Acquisitions (GRAPPA; Griswold et al., 2002) acceleration factor 2; flip angle, 90°; slice matrix size, 64×64; slice thickness, 3.3 mm; slice gap, 0.3 mm; FOV, 212 x 212 mm; bandwidth: 1776 Hz/px; echo spacing: 0.65 ms). A structural image (1 mm isotropic) was acquired using a T1-weighted 3D magnetization-prepared rapid gradient-echo sequence (MP-RAGE; TR, 2.73 s; TE, 2.95 ms; flip angle, 7°; FOV, 256 x 256 x 176 mm).

#### MRI data preprocessing and statistical analyses

MRI data were pre-processed in standard stereotactic (MNI152) space for the purpose of whole-brain group analyses. Mutual information maximization-based rigid-body registration was used to register structural and functional images. Functional images were motion corrected using rigid-body transformations. Structural images were segmented into gray matter, white matter, and CSF images using a unified probabilistic template registration and tissue classification method (Ashburner and Friston, 2005). Tissue images were then registered with site-specific tissue templates (created from 384 T1-weighted scans) using DARTEL (Ashburner, 2007), and registered (using an affine transformation) with the MNI152 template included in SPM8. Identical transformations were applied to all functional images, which were resliced into 2 mm isotropic voxels and smoothed with a 6 mm FWHM Gaussian kernel.

Responses to the two-back task and lateral eye movements were modeled using box-car regressors (duration of 27 s). These two regressors were temporally convolved with the canonical hemodynamic response function (HRF) included in SPM8. Additionally, six movement parameter regressors (3 translations, 3 rotations) derived from rigid-body motion correction, high-pass filtering (1/128 Hz cut-off), and AR(1) serial correlation corrections were included in the model. Single-subject contrast maps of the two-back and eye-movement blocks against fixation were entered into second-level one-sample t-tests.

Finally, we conducted a psychophysiological interaction (PPI) analysis with the amygdala (left and right separately) as seeds for both the eye-movement condition and the two-back condition. We performed the PPI analysis in Experiment 1, because here we used a block design, which is optimal for investigating task-driven connectivity changes on top of task activation (Friston et al., 1997). These (four) first level models were identical to the model described above, but each included two additional regressors namely [1] the timeseries for the first eigenvariate of the amygdala seed (either left or right), and [2] the product of this timeseries [after hemodynamic response function (HRF) deconvolution] with task regressor (either the two-back blocks or the eye movement blocks versus fixation) that was temporally convolved with the canonical HRF included in SPM8. Single-subject contrast maps of the two-back and eye-movement blocks against fixation were entered into second-level full factorial repeated measures ANOVAs with hemisphere as a within-subject variable.

Based on our a priori hypotheses, results for the amygdala and ventral medial prefrontal cortex (vmPFC) were corrected for reduced search volumes using small volume corrections (SVC) and were family-wise error (FWE) corrected using voxel-level statistics. SVC of the amygdala was based on a group mask that was created by averaging individual amygdala segmentations (n=24) of T1-weighted images (using FSL FIRST, see http://www.fmrib.ox.ac.uk/fsl/first/index.html), which were warped into MNI space using DARTEL. The vmPFC was defined as 10 mm sphere around the 0 40 −3 coordinate based on a previous study (Schiller and Delgado, 2010).

#### Eye tracking

For 15 participants, eye tracking was recorded using an MR-compatible eye-tracking system (MEye Track-LR camera unit, SMI, SensoMotoric Instruments). Data were preprocessed using in-house software (Hermans et al., 2013) implemented in Matlab 7.14 (MathWorks). Blinks were removed from the signal using linear interpolation. Eye-tracking data were normalized based on a calibration at the start of the experiment. Analysis revealed participants complied with the instructions of the task.

### EXPERIMENT 2

#### Participants

Twenty-four right-handed healthy volunteers (12 females, 12 males; 20-34 years [M=24.8, SD=3.6]) completed the study. An additional 5 participants did not complete the entire experiment due non-compliance with instructions (*e.g.,* falling asleep). Exclusion criteria were: current or lifetime history of psychiatric, neurological, or endocrine illness, current treatment with any medication that affects central nervous system or endocrine systems, average use of more than 3 alcoholic beverages daily, average use of recreational drugs weekly or more, habitual smoking, predominant left-handedness, uncorrected vision, intense daily physical exercise, and any contraindications for MRI. Participants gave written informed consent and were paid for their participation. This study was approved by the local ethical review board (CMO region Arnhem-Nijmegen).

#### Experimental design

Participants were tested in a differential delay fear conditioning paradigm (Schiller et al., 2010, 2013) on three consecutive days with 24h in between. The first day comprised an acquisition session, the second day an extinction session, and the third day a recall session. The stimulus set across the three days consisted of four squares as conditioned stimuli (CS) with a different color. The luminance of the stimuli, background, and ISI screen was equalized. On day one, two cues (CS+s, 4s duration) were partially reinforced (37.5 % reinforcement rate) with a mild electrical shock to the fingers (*i.e.*, the unconditioned stimulus; UCS). The two other cues (CS-s, 4s duration) were never reinforced. In total, there were 64 trials (16 trials per CS). The CS+s reinforced, CS+s unreinforced, and CS-s were presented in a pseudorandom order. The ISI was jittered between 4s and 8s with an average of 6s.

On day two, extinction included 48 CS trials (12 trials per CS, 4s duration) and 24 eye-movement blocks (10 s duration). One CS+ (CS+_eye_) and one CS-(CS-_eye_) were always followed by an eye-movement block while the other CS+ (CS+_no-eye_) and the other CS-(CS-_no-eye_) were always followed by a fixation block. The ISI between CS and eye movement block was jittered between 0.5s and 1.5s which was done to minimize eye-movement anticipation during the CS presentation. With the duration of 10s, we stayed on the lower end of what is used in EMDR treatment, in which the duration of eye movements varies between 8 and 96 s (Lee and Cuijpers, 2013). This 10 s duration limits the length of the experiment, while still including the peak of the Blood Oxygenation Level Dependent (BOLD) response within the eye-movement blocks (Heeger and Ress, 2002). As in experiment 1, the speed of the moving dot was ∼1 Hz, based on previous laboratory models of EMDR (van den Hout et al., 2013). The visual angle was approximately 11°. We verified compliance of participants using eye-tracking measurements (all participants complied; see below for methodological details, and Figure 4C for averaged results). The ISI after the eye-movement block varied between 4s and 8s with an average of 6s. On day three, the experiment started with a re-extinction session (re-extinction1), which included 24 CS trials (6 trials per CS, 4s duration) with an ISI jittered between 4s and 8s (average of 6s). After this session there was a reinstatement procedure (Haaker et al., 2014) consisting of 3 un-signaled UCS presentations (ISI: 10s). Following this, participants underwent a second re-extinction session (re-extinction2), which included 24 CS trials (6 trials per CS, 4s duration). ISI was jittered between 4s and 8s with an average of 6s. See Figure 2 for an overview.

#### Questionnaires and debriefing

Participants completed the Beck Depression Inventory (BDI; Beck et al., 1996) and the trait version of State-Trait Anxiety Inventory (STAI-t; Van der Ploeg, 1980). A BDI score above 13 was used to exclude participants from the analyses, but none of the participants had a score higher than the cut-off. Average BDI score was 3.5 (range: 0-10) and STAI-t was 33.5 (range: 25-48). Participants were debriefed after the completion of the experiment and asked about their contingency knowledge on the occurrence of electrical shocks, as well as the relationship between the CSs and eye-movement blocks. Participants were furthermore asked about their knowledge of EMDR and whether they at some time during the experiment thought of the experiment in the context of EMDR treatment. Five participants reported doing so. We therefore redid the analyses of the two re-extinction phases on day 3 excluding these five participants. The results and conclusions remained the same and therefore the results are reported including all participants.

#### Peripheral stimulation

Electrical shocks were delivered via two Ag/AgCl electrodes attached to the distal phalanges of the second and third fingers of the right or left hand (counterbalanced between subjects) using a MAXTENS 2000 (Bio-Protech) device. Shock duration was 200 ms, and intensity varied in 10 intensity steps between 0V-40V/0mA-80mA. During a standardized shock intensity adjustment procedure, each participant received and subjectively rated five shocks, allowing shock intensity to converge to a level experienced as uncomfortable, but not painful. The resulting average intensity step was 4.8 (SD: 1.8) on a scale from 1 to 10. The intensity step was set on day 1 and remained the same on day 3 for the reinstatement procedure.

#### Peripheral measurements

Electrodermal activity was assessed using two Ag/AgCl electrodes attached to the distal phalanges of the first and second fingers of the left or right hand (counterbalanced between subjects) using a BrainAmp MR system and recorded using BrainVision Recorder software (Brain Products GmbH, Munich, Germany). Data were preprocessed using in-house software; radio frequency (RF) artifacts were removed and a low-pass filter was applied. Skin conductance responses (SCR) were automatically scored with additional manual supervision using Autonomate (Green et al., 2014) implemented in Matlab 7.14 (MathWorks). We opted to use the magnitude method, since it has been considered the standard method of scoring SCRs for several decades (Edelberg, 1972). SCR amplitudes (measured in μSiem) were determined for each trial within an onset latency window between 0.5 and 4.5 s after stimulus onset, with a minimum rise time of 0.5 s and a maximum rise time of 5 s after response onset. Reinforced trials were omitted and all other response amplitudes were square-root transformed prior to statistical analysis. One subject was omitted from the SCR analyses on day 1 because of failed recordings presumably due to motion of the hand. Analyses on the SCR were performed using SPSS 19 (IBM Corp, Armonk, New York). Four repeated-measures ANOVAs were conducted, one for each experimental phase (Acquisition, Extinction, Re-extinction1, and Re-extinction2). Each ANOVA included CS (CS+, CS-) and Extinction manipulation (eye, no-eye) as within-subject factors. During the extinction and re-extinction phases, an additional within-subject factor was included, namely Time (early, late). Subsequently, differential SCR were calculated (CS+ minus CS-) to test for differences between the two conditions (eye movement and no-eye movement). To test for spontaneous recovery of fear, the differential response on the last trial of extinction was subtracted from the first differential response during re-extinction1 (Schiller et al., 2010, 2013). The reinstatement recovery index was calculated in a similar way by subtracting the last differential response during re-extinction1 from the first differential response during re-extinction2 (Schiller et al., 2010, 2013). For the spontaneous recovery index and reinstatement recovery index analyses, we covaried the order of the CS+ (CS+_eye_ or CS+_no-eye_) presentation. Lastly, the amount of amygdala suppression that occurred on day 2 during the eye-movement blocks was added as a covariate to the recovery index analyses on day 3 to test whether amygdala deactivation predicted fear recovery.

Eye tracking was recorded using an MR-compatible eye-tracking system (MEye Track-LR camera unit, SMI, SensoMotoric Instruments). Data were preprocessed using in-house software (Hermans et al., 2013) implemented in Matlab 7.14 (MathWorks). Blinks were removed from the signal using linear interpolation. Eye-tracking data during the eye-movements blocks were normalized based on a calibration at the start of the experiment.

#### Computational modeling

To further characterize potential differences in the process of extinction learning, we used the computational model from Gershman and Niv (2012) and applied this to acquisition and extinction trials for the eye-movement manipulation and no-eye movement manipulation separately. As input for the model fit we used the square-root transformed SCRs. On each trial, the model assumes that participants determine the probability that a given latent cause generated the observed contingency. The model computes for each participant the preference of assigning all acquisition and extinction trials to a single state or assigning acquisition trials to one state and extinction trials to another state. We performed a comparison between three models, namely “single latent cause” and two models with multiple latent causes, “multiple latent causes without stickiness”, and “multiple latent causes with stickiness”. The stickiness parameter encourages modes in the distribution of states to persist over time (Gershman et al., 2014). To assess the difference between the two extinction manipulations, we first investigated model probabilities over the whole group and next the model preference for each individual. We counted the number of participants per model and tested for a difference in distribution. To test whether the model preference would predict fear recovery, we used the Bayesian Information criterion (BIC) of the models to calculate for each individual a Bayes Factor (BF) which is a ratio of the model preference. A value above 1 means that behavioral evidence favors the multiple latent cause model (without stickiness), and a value below 1 means the evidence favors instead the single latent cause model.

#### Physiological noise correction

Finger pulse was recorded using a pulse oximeter affixed to the third finger of the left or right hand (counterbalanced between subjects). Respiration was measured using a respiration belt placed around the participant’s abdomen. Pulse and respiration measures were used for retrospective image-based correction (RETROICOR) of physiological noise artifacts in BOLD-fMRI data (Glover et al., 2000). Raw pulse and respiratory data were processed offline using in-house software for interactive visual artifact correction and peak detection, and were used to specify fifth-order Fourier models of the cardiac and respiratory phase-related modulation of the BOLD signal (Van Buuren et al., 2009), yielding 10 nuisance regressors for cardiac noise and 10 for respiratory noise. Additional regressors were calculated for heart rate frequency, heart rate variability, (raw) abdominal circumference, respiratory frequency, respiratory amplitude, and respiration volume per unit time (Birn et al., 2006), yielding a total of 26 RETROICOR regressors.

#### MRI data acquisition and multi-echo weighting

MRI scans were acquired using a Siemens (Erlangen, Germany) MAGNETOM Avanto 1.5T MR scanner. T2*-weighted blood oxygenation level-dependent (BOLD) images were recorded using a customized multi-echo EPI sequence with ascending slice acquisition (35 axial slices; TR, 2.2 s; TE, 9.4 ms, 21 ms, 33 ms, 44 ms, and 56 ms; Generalized Autocalibrating Partially Parallel Acquisitions (GRAPPA; Griswold et al., 2002) acceleration factor 3; flip angle, 90°; slice matrix size, 64×64; slice thickness, 3.0 mm; slice gap, 0.51 mm; FOV, 212 x 212 mm; bandwidth: 2604 Hz/px; echo spacing: 0.49 ms). To account for regional variation in susceptibility-induced signal dropout, voxel-wise weighted sums of all echoes were calculated based on local contrast-to-noise ratio (Poser et al., 2006). A structural image (1 mm isotropic) was acquired using a T1-weighted 3D magnetization-prepared rapid gradient-echo sequence (MP-RAGE; TR, 2.73 s; TE, 2.95 ms; flip angle, 7°; FOV, 256 x 256 x 176 mm).

#### MRI data preprocessing in standard stereotactic space and statistical analyses

MRI data were pre-processed in standard stereotactic (MNI152) space for the purpose of whole-brain group analyses. Mutual information maximization-based rigid-body registration was used to register structural and functional images. Functional images were motion corrected using rigid-body transformations. Structural images were segmented into gray matter, white matter, and CSF images using a unified probabilistic template registration and tissue classification method (Ashburner and Friston, 2005). Tissue images were then registered with site-specific tissue templates using DARTEL (Ashburner, 2007), and registered (using an affine transformation) with the MNI152 template included in SPM8. Identical transformations were applied to all functional images, which were resliced into 2 mm isotropic voxels and smoothed with a 6 mm FWHM Gaussian kernel.

We created four first-level models for all stages of the experiment (i.e., Acquisition, Extinction, Re-ectinction1 and Re-extinction2). Responses to the CSs were modeled using box-car regressors (duration of 5s). During the acquisition phase, additional regressors included the reinforced CS+s (duration of 5s) and the shock which was modeled as a stick function. During the extinction phase, responses to the eye-movement blocks were modeled using box-car regressors with a duration of 10s. Regressors were temporally convolved with the canonical hemodynamic response function (HRF) included in SPM8. Additionally, six movement parameter regressors (3 translations, 3 rotations) derived from rigid-body motion correction, high-pass filtering (1/128 Hz cut-off), and AR(1) serial correlation corrections were included in the model. Single-subject contrast maps were entered into second-level one-sample t-tests.

Although a PPI analysis in experiments with event-related designs is more difficult to interpret (O’Reilly et al., 2012), we did perform a PPI analysis, similar to Experiment 1, with the amygdala as a seed region in Experiment 2 as well. We did not find any statistically significant connectivity changes. As an extra check, we used the FEF as a seed as well, but also here we did not observe connectivity differences. There are two possible explanations for this. First, the shape and assumptions regarding the HRF are more important in event-related designs than in blocked designs (Gitelman et al., 2003). Second, PPI analyses have less power than task activity analyses, and event-related designs tend to have smaller effect sizes than block designs. This is because the period where it is possible to look at task-driven connectivity changes are shorter than in block designs.

Based on our priori hypotheses, results for the amygdala and dorsal anterior cingulate cortex (dACC) were corrected for reduced search volumes using small volume corrections (SVC) and were family-wise error (FWE) corrected using voxel-level statistics. SVC of the amygdala was based on a group mask that was created by averaging individual amygdala segmentations (n=24) of T1-weighted images (using FSL FIRST, see http://www.fmrib.ox.ac.uk/fsl/first/index.html), which were warped into MNI space using DARTEL. The dACC was defined based on a functional ROI atlas (Shirer et al., 2012).

#### MRI data preprocessing of the extinction session in native space and statistical analyses

For the primary fMRI analysis (amygdala response during eye movements), we preprocessed MRI data during extinction in native space (*i.e.*, without stereotactic normalization) using SPM8 (http://www.fil.ion.ucl.ac.uk/spm; Wellcome Department of Imaging Neuroscience, London, UK). Since the results from Experiment 1 showed the suppression was not specific to the amygdala per se, we opted for this more specific analysis to make sure that the effects in the amygdala are not, for example, due to signal blurring from the hippocampus into the amygdala. Additionally, this analysis allowed us to extract an averaged time course of the amygdala signal.

All functional scans were co-registered with structural scans using mutual information maximization. The amygdala was individually defined in native space using automated anatomical segmentation of T1-weighted images using FSL FIRST (see http://www.fmrib.ox.ac.uk/fsl/first/index.html). The amygdala segmentations underwent visual inspection. We observed that some amygdala voxels for some participants were not part of the mask, indicating that the FSL FIRST segmentation was relatively conservative (*e.g.,* in anterior parts, which was depicted in Moore et al., 2014, as well). For each participant, we made sure that the amygdala mask did not contain voxels that were part of, or extended into, the hippocampus. We therefore conclude that our amygdala masks are representative of amygdala volume, and that the suppression effect can be assigned to the amygdala. The ventral medial prefrontal cortex (vmPFC) was defined as 10 mm sphere around the 0 40 −3 coordinate based on a previous study (Schiller and Delgado, 2010). The Frontal Eye Fields (FEF) were defined based on a 5 mm sphere around the MNI peak coordinates reported in a meta-analysis (Jamadar et al., 2013). Subsequently, the FEF masks were transferred back into native space for each individual using the reversed spatial normalization parameters.

For statistical analyses, responses to the eye-movement and no-eye movement blocks were estimated using a finite impulse response (FIR) model which included 9 time bins (TR = 2.2 s) starting one time-bin before the onset of the CS (−2.2 s) and ending one time bin after the eye-movement blocks (17.6 s). Therefore, bin numbers 5 until 8 (6.6 s – 15.4 s) always fell within the eye-movement blocks. For the no-eye movement block, the same time frame was used. This first-level model makes no assumptions regarding the HRF shape, and yields independent response estimates for all 9 time bins, which makes it possible to investigate the time course of the responses. The first-level models additionally included six movement parameter regressors (3 translations, 3 rotations) derived from rigid-body motion correction, 26 RETROICOR physiological noise regressors (see above), high-pass filtering (1/128 Hz cut-off), and AR(1) serial correlations correction. We extracted the average beta weights within the amygdala and FEF for each time bin and each CS. A repeated-measures ANOVA was conducted for each region with CS (CS+, CS-), Extinction manipulation (eye, no-eye) and Time bin (5-8) as within-subject factors. See Figure 4.

#### Sample size calculation, statistical testing, and data sharing

Sample size of 24 was determined based a pooled effect size from four studies that have used similar paradigms (Schiller et al., 2010, 2013; Agren et al., 2012; Kindt and Soeter, 2013), alpha=.05, and 1-beta=.80. Partial eta squared (*Pη²*) or Cohen’s d effect size estimates are reported for all relevant tests. Alpha was set at .05 throughout and two-tailed t-tests were conducted unless stated otherwise. Anonymized data will be made available to others upon formal request.

## Results

### EXPERIMENT 1

#### A working memory task and goal-directed eye movements suppress amygdala activity

In a block design, participants performed a two-back working memory task and goal-directed eye movements while undergoing functional MRI. We found typical activation patterns within the dorsal fronto-parietal network (Corbetta and Shulman, 2002; Qin et al., 2009) during the two-back [*e.g*., the left and right dorsolateral PFC: *p*<.001 peak-voxel FWE-SVC, and left and right posterior parietal cortex: *p*<.001 peak-voxel FWE-SVC] and eye-movement blocks [*e.g*., the left and right frontal eye fields: *p*<.001 peak-voxel FWE-SVC, and left and right posterior parietal cortex: *p*<.001 peak-voxel FWE-SVC]. Further, and as expected, both the two-back blocks [left: *p*=.001, right: *p*=.003; peak-voxel FWE-SVC] and eye-movement blocks [left: *p*=.035; right: *p*=.08, peak-voxel FWE-SVC] led to deactivations in the amygdala compared to fixation. The amygdala suppression during the eye-movement blocks was not as strong (*i.e.*, suppression was only significant in the left amygdala), however a direct comparison revealed no difference in amygdala deactivation between the two-back and eye-movements blocks. When using the two-back blocks as a functional localizer for the amygdala, the suppression was significant as well [left: *p*=.015; right: *p*=.044, peak-voxel FWE-SVC], indicating the suppression is in a similar location for both tasks. Among other regions, deactivation was also found in the ventromedial PFC (*p*<.001, FWE-SVC) in the two-back blocks. For all whole-brain activation and deactivation results see Figure 1B and Table 1. See Figure 1D for an illustration of the location of the suppression in the amygdala.

**Figure 1.**
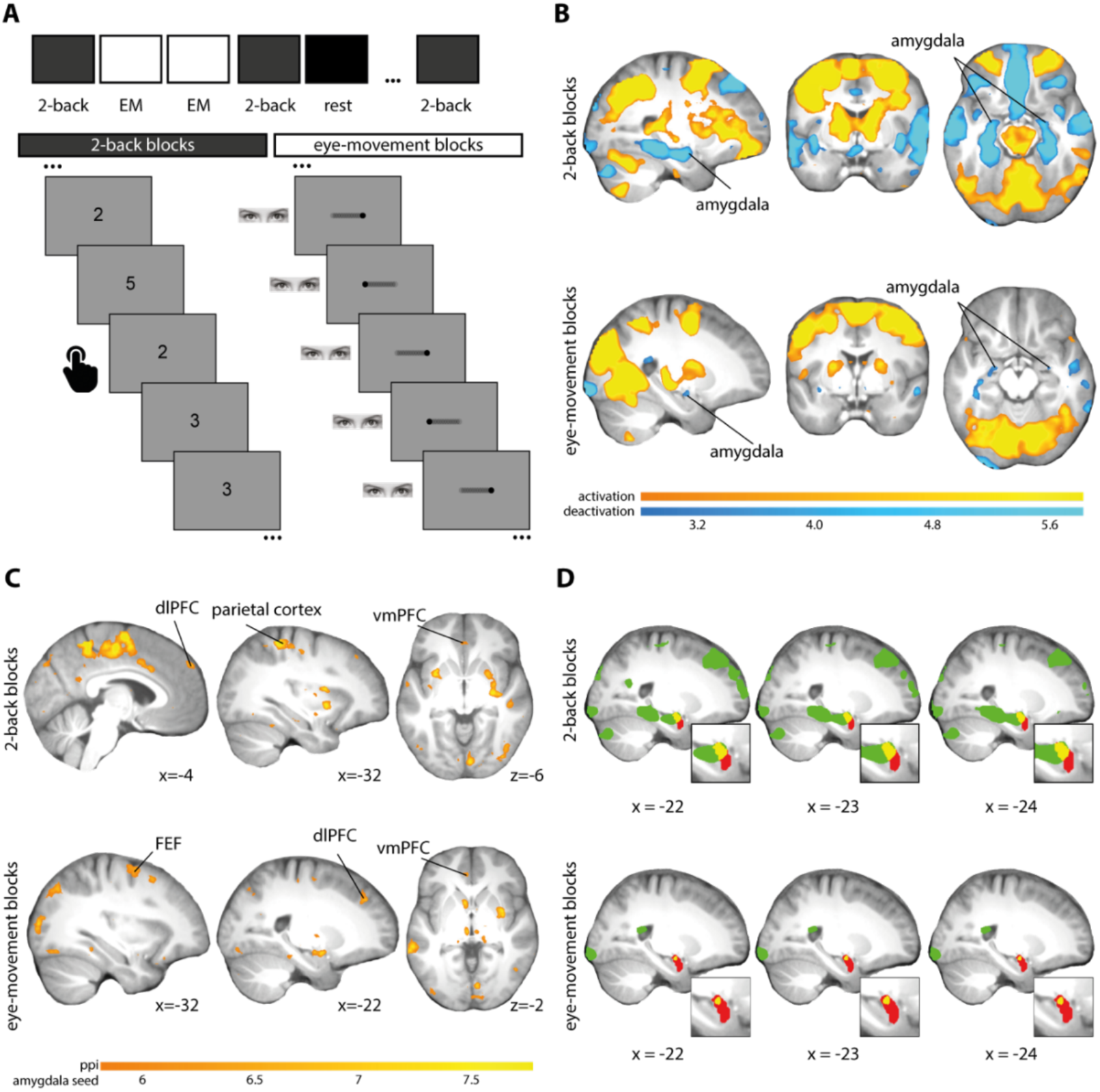
(**A**) Experimental design. (**B**) Activations and deactivations of the two-back task and eye movements compared to fixation. The threshold for significance for the amygdala (whole-brain threshold p<.005 uncorrected and peak voxel FWE-SVC p<.05) is applied to the whole brain to show the specificity of the effect. Whole-brain corrected inferential statistics are reported in Table 1. (**C**) PPI cluster with the left and right amygdala as a seed for the two-back task and eye movements compared to fixation. Images are thresholded at p<.05, whole-brain FWE corrected. (**D**) Deactivation during the two-back blocks and eye-movement blocks (in green) overlaid onto the average T1 scan from all participants. In red is the anatomical location of the amygdala. It can be seen that the deactivation (overlap is in yellow) is located towards the dorsal part in both tasks. EM: eye movements.

**Table 1.** Peak voxel coordinates and statistics of activations and deactivations of the two-back task and eye movements compared to fixation (Experiment 1)

| Region | Side | x(mm) | y(mm) | z(mm) | Peak T | P-value |
| --- | --- | --- | --- | --- | --- | --- |
| *Two-Back blocks > fixation <sup>a</sup>* |  |  |  |  |  |  |
| Anterior insula | R | 34 | 26 | 0 | 14.70 | P<.001 |
| Dorsolateral prefrontal cortex (dlPFC) | L | -40 | 2 | 30 | 8.57 | P<.001 |
| Dorsolateral prefrontal cortex (dlPFC) | R | 44 | 38 | 24 | 7.16 | P<.001 |
| Posterior parietal cortex | L | -34 | -56 | 50 | 11.63 | P<.001 |
| Posterior parietal cortex | R | 48 | -36 | 46 | 11.88 | P<.001 |
| *Two-Back blocks < fixation* |  |  |  |  |  |  |
| Superior frontal gyrus | L | -18 | 42 | 46 | 13.38 | P<.001 |
| Precuneus | L | -4 | -52 | 14 | 12.72 | P<.001 |
| Cerebellum | R | 34 | -78 | -34 | 10.80 | P<.001 |
| Rolandic operculum | R | 48 | -22 | 18 | 10.12 | P<.001 |
| Angular gyrus | L | -48 | -64 | 26 | 10.08 | P<.001 |
| Middle temporal gyrus | L | -58 | -8 | -6 | 9.69 | P<.001 |
| Fusiform gyrus | L | -32 | -40 | -12 | 9.60 | P<.001 |
| Angular gyrus | R | 56 | -66 | 36 | 9.25 | P<.001 |
| Fusiform gyrus | R | 30 | -42 | -10 | 7.83 | P=.007 |
| Inferior frontal gyrus | R | 52 | 32 | 6 | 7.80 | P=.007 |
| Inferior orbital frontal gyrus | R | 38 | 36 | -10 | 7.44 | P=.015 |
| Cerebellum | L | -26 | -84 | -32 | 7.13 | P=.025 |
| Ventral medial prefrontal cortex (vmPFC) | L/R | -6 | 46 | 0 | 8.39 | P<.001* |
| Amygdala | L | -26 | -10 | -18 | 5.64 | P=.001* |
| Amygdala | R | 24 | -8 | -16 | 4.73 | P=.003* |
| *Eye movements > fixation* |  |  |  |  |  |  |
| Calcarine | R | 6 | -70 | 8 | 7.37 | P<.001 |
| Precentral gyrus (Frontal Eye Fields) | L | -40 | -10 | 50 | 6.57 | P<.001 |
| Precentral gyrus (Frontal Eye Fields) | R | 32 | -4 | 52 | 8.48 | P=.002 |
| Posterior parietal cortex | R | 28 | -52 | 50 | 5.81 | P=.001 |
| Posterior parietal cortex | L | -26 | -52 | 54 | 5.77 | P=.001 |
| Middle cingulate cortex | L | -12 | -20 | 40 | 5.22 | P=.018 |
| *Eye movements < fixation* |  |  |  |  |  |  |
| Amygdala | L | -24 | -8 | -14 | 3.17 | P=.035* |
| Amygdala | R | 28 | -8 | -16 | 3.13 | P=.08* |
| Inferior occipital gyrus | R | 34 | -98 | -8 | 5.38 | P=.008 |
| Inferior occipital gyrus | L | -26 | -98 | -10 | 5.31 | P=.012 |

Notes: all coordinates are defined in MNI152 space. All statistics listed are significant at $p<.05$ , whole-brain family-wise error corrected unless indicated otherwise. <sup>a</sup>To reduce the number of peak voxels, only the peak voxels in the predefined ROIs are reported. \*Small volume corrected for region of interest.

Finally, we conducted a psychophysiological interaction (PPI) analysis and found enhanced coupling between the amygdala and the dorsal fronto-parietal network during both the two-back [e.g., the left and right dorsolateral PFC: *p*<.001 peak-voxel FWE-SVC] and eye-movement blocks [e.g., the left and right dorsolateral PFC: *p*<.001 peak-voxel FWE-SVC, and left and right frontal eye fields: p<.001 peak-voxel FWE-SVC]. We also found enhanced coupling between the amygdala and ventromedial PFC, during both the two-back [*p*<.001 peak-voxel FWE-SVC] and eye-movement blocks [*p*=.001 peak-voxel FWE-SVC]. For all connectivity results see Figure 1C and Table 2.

**Table 2.** Peak voxel coordinates and statistics of PPI analyses (with the amygdala as a seed) of the two-back task and eye movements compared to fixation (Experiment 1)

| Region | Side | x(mm) | y(mm) | z(mm) | Peak T | P-value |
| --- | --- | --- | --- | --- | --- | --- |
| *Two-Back blocks > fixation* |  |  |  |  |  |  |
| Superior motor area (SMA) | R | 2 | -4 | 60 | 11.10 | p<.001 |
| Dorsolateral prefrontal cortex (dlPFC) | L | -4 | 56 | 36 | 6.84 | P<.001 |
| Dorsolateral prefrontal cortex (dlPFC) | R | 62 | 6 | 24 | 6.72 | P<.001 |
| Posterior parietal cortex | L | -32 | -40 | 56 | 8.63 | P<.001 |
| Posterior parietal cortex | R | 6 | -36 | 48 | 7.36 | P<.001 |
| Ventromedial prefrontal cortex (vmPFC) | L/R | -2 | 38 | -6 | 6.12 | P<.001* |
| *Eye movements > fixation* |  |  |  |  |  |  |
| Middle cingulate cortex | R | 6 | -38 | 34 | 10.21 | P<.001 |
| Dorsolateral prefrontal cortex (dlPFC) | L | -22 | 40 | 36 | 6.89 | P<.001 |
| Dorsolateral prefrontal cortex (dlPFC) | R | 4 | 66 | 10 | 6.52 | P<.001 |
| Posterior parietal cortex | L | -4 | -62 | 30 | 7.47 | P<.001 |
| Posterior parietal cortex | R | 52 | -24 | 14 | 7.24 | P<.001 |
| Precentral gyrus (frontal eye fields) | L | -44 | -18 | 46 | 7.01 | P<.001 |
| Precentral gyrus (frontal eye fields) | R | 44 | -14 | 56 | 7.04 | P<.001 |
| Ventromedial prefrontal cortex (vmPFC) | L/R | -4 | 42 | -2 | 6.44 | P<.001* |

Notes: all coordinates are defined in MNI152 space. All statistics listed are significant at $p<.05$ , whole-brain family-wise error corrected unless indicated otherwise. To reduce the number of peak voxels, only the peak voxels in the predefined ROIs are reported. \*Small volume corrected for region of interest

In conclusion, goal-directed eye movements, similar to a working memory task (Qin et al., 2009), suppress amygdala activity and induce enhanced coupling between the amygdala and regions involved in cognitive regulation of emotion (Ochsner and Gross, 2005).

### EXPERIMENT 2

#### Goal-directed eye movements suppress amygdala activity

First, we investigated amygdala activity in response to the goal-directed eye movements during extinction learning. We analyzed the data in native space to make sure that the effects in the amygdala and not due to signal blurring from the hippocampus into the amygdala.

Replicating Experiment 1, goal-directed eye movements increased frontal eye field activity [*F*(1,23)=13.11, *p*=.001, *Pη2*=.36] compared to fixation. Amygdala activity was decreased [*F*(1,23)=4.576, *p*=.04, *Pη^2^*=.17] compared to fixation and there was no interaction with CS (CS+, CS-) [*F*(1,23)=1.296, *p*=.27, *Pη^2^*=.05].

Thus, in two independent studies we found that goal-directed eye movements suppress amygdala activity. See Figure 4B.

#### Skin conductance responses during fear acquisition and extinction

SCR measures during acquisition (Figure 3A) revealed a robust differential conditioning effect (CS+ versus CS-) across all trials [*F*(1,22)=18.54, *p*=2.86E-4, *Pη^2^*=.46] and there was no interaction with later Extinction manipulation (eye, no-eye) [*F*(1,22)=1.945, *p*=.18, *Pη^2^*=.08]. During early extinction, there was a differential conditioning effect [*F*(1,23)=49.77, *p*=3.46E-7, *Pη^2^*=.68] which became non-significant during late extinction [*F*(1,23)=.896, *p*=.35, *Pη^2^*=.04] (Figure 3B). There was full extinction on the last trial [*F*(1,23)=.260, *p*=.61, *Pη^2^*=.01] and no interaction with Extinction manipulation (eye, o-eye) [*F*(1,23)=.991, *p*=.33, *Pη^2^*=.04]. Thus, SCR measures revealed there was successful acquisition and extinction, which did not differ significantly between the eye movement and no-eye movement manipulation.

**Figure 2.**
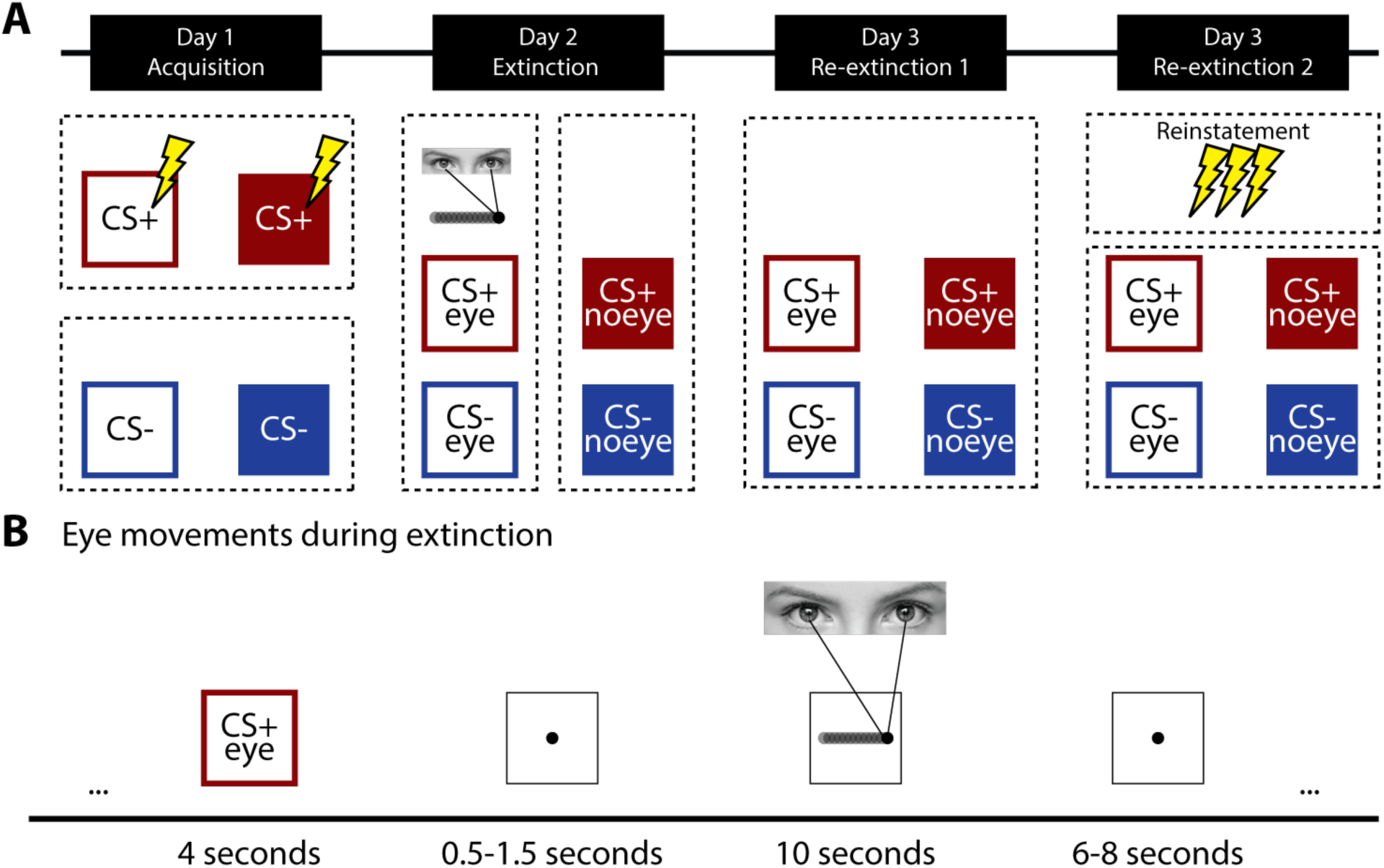
Overview of the experimental design. **(A)** The experiment took place on three consecutive days and included an acquisition phase (day one), an extinction phase (day two), and a re-extinction (re-extinction1) phase and re-extinction after reinstatement (re-extinction2) phase (day three). **(B)** Time line of a single trial during extinction when participants made eye movements. CS: conditioned stimulus.

**Figure 3.**
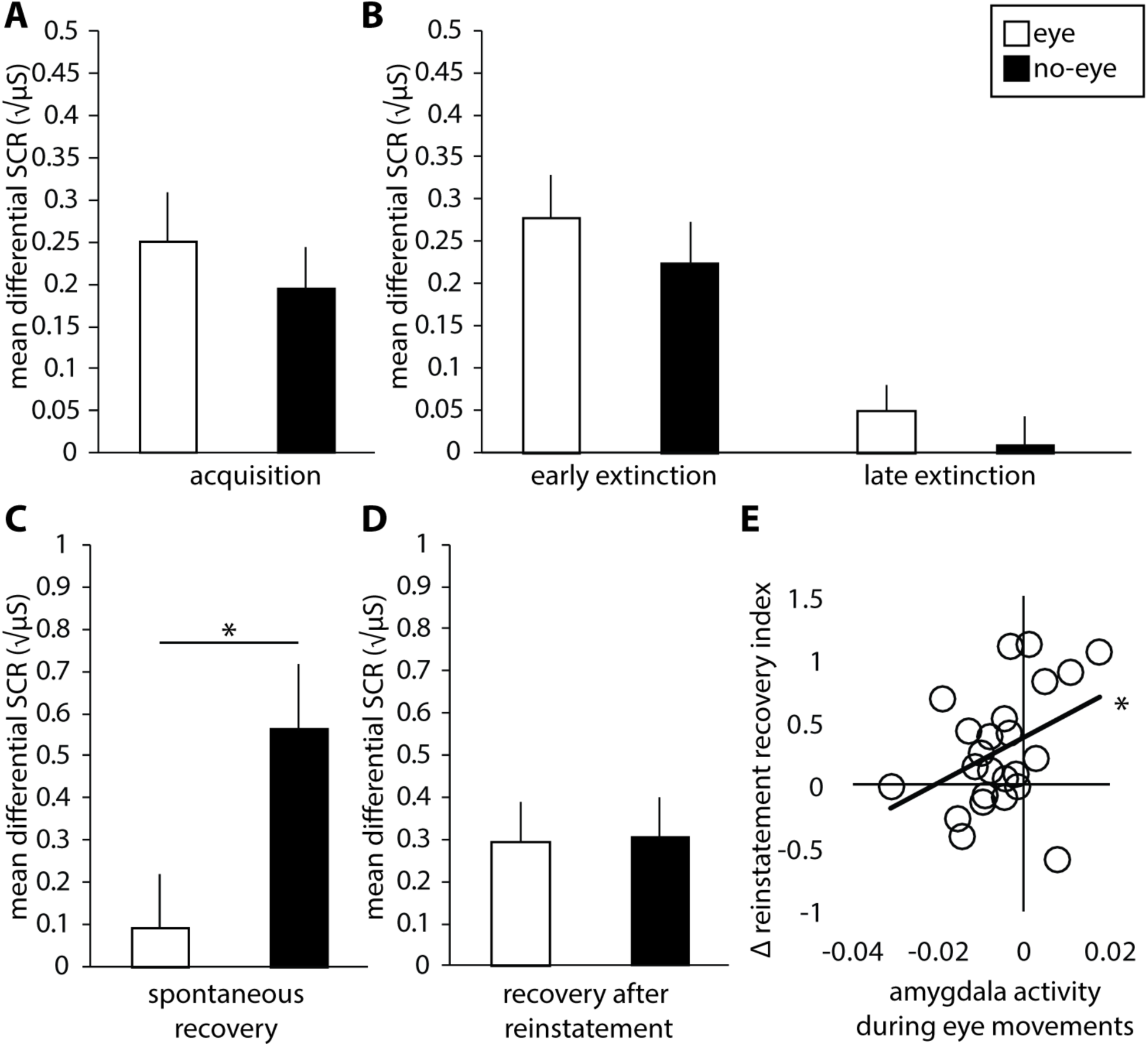
Differential skin conductance responses (SCR) measured during (**A**) acquisition and (**B**) early and late extinction. (**C**) Differential spontaneous recovery index (first trial of re-extinction1 minus last trial of extinction) and (**D**) differential reinstatement recovery index (first trial re-extinction2 minus last trial re-extinction1). (**E**) Correlation between amygdala deactivation during eye-movement blocks on day two and differential reinstatement recovery index for extinction learning with eye movements. Error bars represent ± standard error of the mean. *= p < .05.

**Figure 4.**
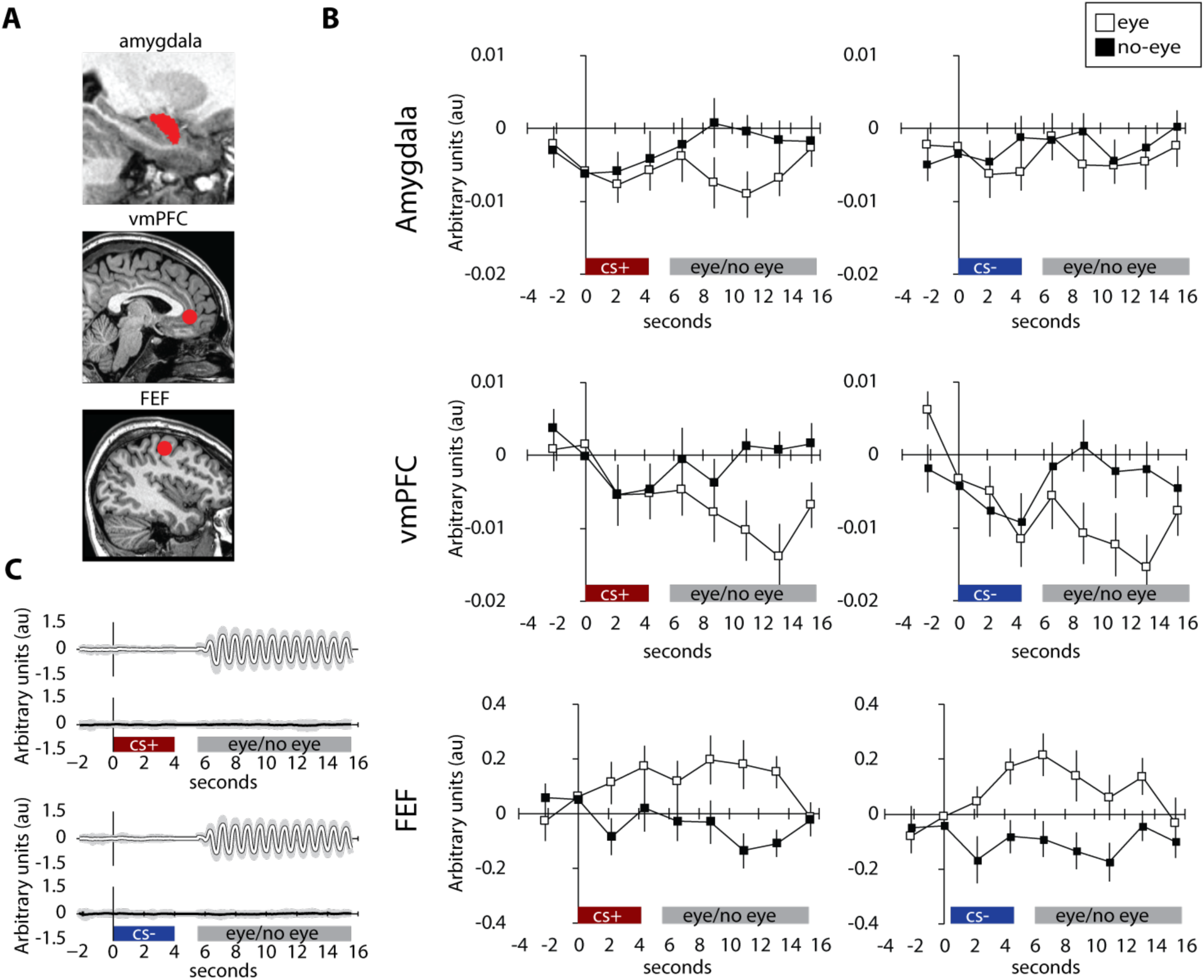
Eye movements suppress amygdala activity. (**A**) Single-subject example of the automated amygdala segmentation, ventral medial prefrontal cortex (vmPFC) was defined by a 10 mm radius around the 0 40 −3 coordinate (Schiller and Delgado, 2010) and frontal eye field (FEF) as a bilateral 5 mm radius around the MNI peak coordinates reported in a meta-analysis of neuroimaging studies of eye movements (Jamadar et al., 2013). (**B**) Amygdala and vmPFC deactivation and FEF activation during eye-movement and no-eye movement blocks within the extinction phase. Error bars represent ± SEM. (**C**) Eye-movement recordings during eye-movement and no-eye movement blocks within the extinction phase. The black and white lines reflect the mean across all participants and the gray shaded area the standard error of the mean (SEM).

#### Eye movements during extinction block spontaneous recovery of fear the following day

Crucially, and as predicted, a repeated-measures ANOVA across all re-extinction1 trials on day three revealed an interaction between Extinction manipulation (eye, no-eye) and Time (first versus second half of re-extinction1) [*F*(1,22)=6.723, *p*=.02, *Pη^2^*=.23]. Follow-up tests on the spontaneous recovery index, indicated by a differential responding from the last trial of extinction to the first trial of re-extinction1 (Schiller et al., 2010), revealed that spontaneous recovery differed between extinction manipulations [*F*(1,22)=5.976, *p*=.02, *Pη^2^*=.21]. As expected, there was spontaneous recovery for extinction without [*t*(22)= 3.60, *p*=.002], but not with [*t*(22)=.694, *p*=.50; Figure 3C] eye movements. Thus, eye movements during extinction learning indeed blocked spontaneous recovery.

#### Eye movement-induced amygdala suppression during extinction predicts a reduction in fear recovery after reinstatement

Analyses on the reinstatement recovery index, indicated by a differential responding from the last trial of re-extinction1 to the first trial of re-extinction2 (Schiller et al., 2010), showed that differential responses returned on average [*F*(1,22)=23.486, *p*=7.65E-5, *Pη^2^*=.52], and that there was no interaction with Extinction manipulation [*F*(1,22)=.005, *p*=.94, *Pη^2^*<.001]. Notably, including strength of amygdala deactivation as a covariate revealed an interaction between this deactivation and Extinction manipulation [*F*(1,21)=7.252, *p*=.01, *Pη^2^*=.26]. Follow-up tests showed a positive correlation between amygdala deactivation and recovery for the eye-movement manipulation [*r*(21)=.39 *p*=.028, one-tailed; Figure 3E]. Amygdala responses during the no-eye movement blocks did not predict recovery following reinstatement [*F*(1,21)=1.392, *p*=.25, *Pη^2^*=.06]. Additionally, for the spontaneous recovery index, this association was not found [*F*(1,21)=1.592, *p*=.22, *Pη^2^*=.07]. However, amygdala deactivation did predict the difference between the spontaneous recovery index and reinstatement recovery index [*r*(20)=.62 *p*=.002]. Responses in the frontal eye fields were not associated with recovery following reinstatement [*F*(1,21)=.44, *p*=.51, *Pη^2^*=.02].

In conclusion, differential fear responses on average recovered after reinstatement, however, recovery for the eye-movement condition was attenuated when participants had stronger amygdala deactivations during eye movements. The observation of recovery of fear following reinstatement speaks against a strong interpretation of our findings as resulting from updating (or unlearning) of the original memory trace, and thus favors an account in terms of enhanced extinction learning (i.e., new learning).

#### Computation modeling of skin conductance responses

To further elucidate the mechanism via which eye movements reduced fear recovery, we additionally applied computational modeling (Gershman and Niv, 2012; Gershman and Hartley, 2015) to the SCR data acquired during extinction learning. We asked whether this analysis would reveal a difference in learning not accounted for when averaging across trials. We reasoned that if the eye movements reduce fear recovery via the process of unlearning, individuals would more often assign acquisition and extinction trials to the same latent cause (*i.e.*, update the original memory trace). By contrast, if the eye movements enhance new learning, then acquisition and extinction trials would more likely arise from multiple latent causes (*i.e.*, forming a safety memory while leaving the original memory trace intact; Gershman and Hartley, 2015).

We compared three models and found that for standard extinction all participants aligned with the “single latent cause” model rather than the two models with “multiple latent causes”, with a probability averaged across participants of .92 over .05. For the eye-movement manipulation we saw that four out of 23 participants aligned with the “multiple latent causes without stickiness” model rather than the “single latent cause” model, with a probability of .81 over .16. The “single latent cause” model was still the winning model over the whole sample as indicated by an average model probability >.80 in both extinction conditions.

Since there was a numerical difference in the number of participants in model preference, we further assessed the relationship between model preference (single state vs. multiple latent states) and extinction manipulation (eye, no-eye). We performed a Chi-square test and found that the distribution is significantly different between the two conditions [χ2=4.381, *p*=.036]. A Bayesian Chi-square test implemented in the JASP software (The JASP Team., 2017; https://jasp-stats.org/) indicated moderate support for the alternative hypothesis (more single state models in the standard extinction manipulation in comparison to the eye-movement manipulation) as indicated by a Bayes Factor (BF) of 4.32 in favor of the alternative.

To test whether the state preference would predict the reduction in fear recovery, we next obtained a ratio of the model preference for each individual by using the Bayesian information criterion (BIC) of both models to calculate the BF for the eye-movement manipulation. A value above 1 means that behavioral evidence favors the multiple latent cause model (*i.e.*, this was the case for the four out of 23 participants mentioned above, for the eye-movement manipulation) and a value below 1 means the evidence instead favors the single latent cause model. Neither the spontaneous recovery index nor the recovery following reinstatement index correlated with the BF.

Thus, overall most individuals in our sample were one-state learners, consistent with previous findings (Gershman and Hartley, 2015). If eye movements would further enhance unlearning, then we would have expected to observe a greater preference for a single latent cause in the eye-movement condition compared to standard extinction. The fact that we observed the opposite (i.e., a stronger preference for multiple latent causes in the eye-movement condition) is therefore also at odds with the notion that eye-movement manipulations would enhance updating or unlearning.

#### Whole-brain group analysis in standard stereotactic space

We first verified whether the acquisition phase exhibited the expected task-related activation and deactivation during CS presentation using conventional group analyses in standard stereotactic (MNI152) space. We observed robust differential BOLD responses in the anterior insula [left: *p*<.001 right: *p*<.001 peak-voxel FWE-corrected] and dorsal anterior cingulate cortex (dACC) [*p*<.001 peak-voxel FWE-corrected] among others. Additionally, we observed robust deactivations in the vmPFC [*p*=.007 peak-voxel FWE-corrected] among others. See Table 3.

**Table 3.** Peak voxel coordinates and statistics during acquisition

| Region | Side | x(mm) | y(mm) | z(mm) | Peak T | P-value |
| --- | --- | --- | --- | --- | --- | --- |
| *CS+ > CS-* |  |  |  |  |  |  |
| Inferior frontal gyrus / anterior insula | R | 50 | 24 | 4 | 9.6797 | P<.001 |
| Inferior frontal gyrus / anterior insula | L | -36 | 24 | 4 | 8.90 | P<.001 |
| Supramarginal gyrus | R | 54 | -40 | 28 | 8.6493 | P<.001 |
| Dorsal anterior cingulate cortex (dACC) | R | 8 | 26 | 28 | 7.8007 | P<.001 |
| Supramarginal gyrus | L | -54 | -40 | 32 | 6.1253 | P=.002 |
| Cerebellum | R | 2 | -54 | -36 | 5.9356 | P=.004 |
| Cerebellum | L | -34 | -54 | -32 | 5.9167 | P=.004 |
| *CS- > CS+* |  |  |  |  |  |  |
| Middle occipital gyrus | L | -44 | -74 | 30 | 5.8497 | P=.006 |
| Hippocampus | L | -24 | -18 | -18 | 5.8302 | P=.006 |
| Ventral medial prefrontal cortex (vmPFC) | L | -10 | 52 | -6 | 5.7878 | P=.007 |
| Precuneus | R | 10 | -54 | 14 | 5.7507 | P=.008 |
| Paracentral lobule | R | 8 | -26 | 70 | 5.6498 | P=.012 |
| Precuneus | L | -8 | -60 | 16 | 5.6123 | P=.014 |
| Fusiform gyrus / Para-hippocampal gyrus | R | 32 | -44 | -4 | 5.3385 | P=.038 |

Notes: all coordinates are defined in MNI152 space. All statistics listed are significant at $p<.05$ , whole-brain family-wise error corrected unless indicated otherwise.

Next, we investigated response patterns to the CS during extinction learning. First, we found activation patterns in the anterior insula [left: *p*=.016 right: *p*=.043 peak-voxel FWE-corrected] and dACC [*p*=.02 peak-voxel FWE-SCV] as a main effect of CS (CS+ versus CS-). We did not observe any deactivations. Interestingly, similar to what our native space analysis of the frontal eye fields (FEF) activation already showed (see Figure 4B), we found activation in the FEF in response to the CS that was coupled to the eye movements [left: *p*=.034 peak-voxel FWE-corrected; right: *p*=.001 peak-voxel FWE-SVC] as a main effect of eye movements. This was before the execution of the eye movements and thus indicates an anticipation response. There was no CS (CS+, CS-) by Extinction manipulation (eye, no-eye) interaction.

Critically, in response to the eye movement blocks (which followed the CS presentation after a brief delay) we found deactivation in the vmPFC [*p*=.025 peak-voxel FWE-corrected]. See Table 4 for the full whole-brain results. Additional analysis on the vmPFC during the native space data confirmed vmPFC was deactivated compared to fixation [*F*(1,23)=7.265, *p*=.013, *Pη2*=.24]. This deactivation did not predict recovery following reinstatement [*F*(1,23)=.065, *p*=.801, *Pη^2^*=.003]. Thus, similar to amygdala responses, vmPFC responses are suppressed during the eye-movement blocks.

**Table 4.** Peak voxel coordinates and statistics during extinction learning

| Region | Side | x(mm) | y(mm) | z(mm) | Peak T | P-value |
| --- | --- | --- | --- | --- | --- | --- |
| **CS presentation** |  |  |  |  |  |  |
| *CS+ > CS- [main effect CS]* |  |  |  |  |  |  |
| Anterior insula | R | 42 | 18 | -6 | 6.36 | P=.001 |
| Anterior insula | L | -32 | 22 | -4 | 5.75 | P=.009 |
| Frontal middle orbital cortex | R | 20 | 60 | -12 | 5.61 | P=.02 |
| Dorsal anterior cingulate cortex (dACC) | R | 4 | 28 | 30 | 5.31 | P=.04 |
| *Eye movements > No-eye movements [main effect eye movements]* |  |  |  |  |  |  |
| Calcrine | L | 10 | -68 | 18 | 10.82 | P<.001 |
| Supplemental Motor Area | L | -4 | -6 | 62 | 7.94 | P<.001 |
| Posterior parietal cortex | L | -26 | -52 | 52 | 7.60 | P<.001 |
| Precentral gyrus (frontal eye fields) | L | -44 | -8 | 52 | 7.58 | P<.001 |
| Precentral gyrus (frontal eye fields) | R | 44 | -8 | 50 | 7.94 | P<.001 |
| Posterior parietal cortex | R | 26 | -52 | 48 | 7.53 | P<.001 |
| Middle temporal gyrus | R | 44 | -64 | 10 | 6.26 | P=.001 |
| Middle cingulate cortex | L | -14 | -20 | 38 | 5.98 | P=.044 |
| **Eye movement blocks** |  |  |  |  |  |  |
| *Eye movements > fixation* |  |  |  |  |  |  |
| Lingual gyrus | L | -10 | -88 | -2 | 15.13 | p<.001 |
| Precentral gyrus (frontal eye fields) | R | 48 | -4 | 42 | 10.26 | P<.001 |
| Putamen | L | -22 | -2 | 12 | 9.80 | P<.001 |
| Supplemental Motor Area | L | -4 | -4 | 62 | 8.21 | P=.003 |
| Precentral gyrus (frontal eye fields) | L | -52 | -4 | 38 | 8.02 | P=.005 |
| Putamen | R | 22 | 8 | 10 | 7.46 | P=.016 |
| *Eye movements < fixation* |  |  |  |  |  |  |
| Posterior insula | R | 38 | -18 | 12 | 11.22 | P<.001 |
| Parahippocampal gyrus | R | 18 | -14 | -24 | 9.16 | P<.001 |
| Posterior insula | L | -36 | -20 | 8 | 8.99 | P=.001 |
| Inferior occipital gyrus | R | 32 | -96 | -4 | 7.80 | P=.008 |
| Superior parietal gyrus | R | 18 | -44 | 68 | 7.44 | P=.017 |
| Inferior occipital gyrus | L | -28 | -94 | -6 | 7.34 | P=.020 |
| Ventral medial prefrontal cortex (vmPFC) | L | -8 | 24 | -12 | 7.22 | P=.025 |
| Post-central gyrus | R | 44 | -26 | 54 | 7.01 | P=.036 |
| Amygdala | R | 32 | 4 | -20 | 4.43 | P=.008* |

Notes: all coordinates are defined in MNI152 space. All statistics listed are significant at $p<.05$ , whole-brain family-wise error corrected unless indicated otherwise.
\*Small volume corrected for region of interest.

**Table 5.** Peak voxel coordinates and statistics during re-extinction1 and re-extinction2

| Region |  | Side | x(mm) | y(mm) | z(mm) | Peak T | P-value |
| --- | --- | --- | --- | --- | --- | --- | --- |
| **Re-extinction1** |  |  |  |  |  |  |  |
| *Cs+ > CS-* |  |  |  |  |  |  |  |
|  | Anterior insula | R | 50 | 20 | -2 | 6.3 | P<.001 |
| **Re-extinction2** |  |  |  |  |  |  |  |
| *CS+ > CS-* |  |  |  |  |  |  |  |
|  | Dorsal Anterior cingulate cortex | R | 8 | 26 | 50 | 5.52 | P=.024 |
|  | Thalamus | R | 6 | -6 | 2 | 5.52 | P=.024 |
| *Cs by Extinction manipulation interaction* |  |  |  |  |  |  |  |
|  | Dorsal Anterior cingulate cortex | L | -8 | 28 | 38 | 4.89 | P=.003* |
| *CS+ > CS- [no-eye movements]* |  |  |  |  |  |  |  |
|  | Dorsal Anterior cingulate cortex | R | 8 | 36 | 40 | 6.24 | P=.007 |

Notes: all coordinates are defined in MNI152 space. All statistics listed are significant at $p < .05$ , whole-brain family-wise error corrected unless indicated otherwise.
\*Small volume corrected for region of interest.

Finally, during re-extinction1, we found differential BOLD responses (CS+ versus CS-) in the anterior insula [*p*<.001 peak-voxel FWE-corrected]. During re-extinction2, we found differential BOLD responses in the dACC [*p*<.001 peak-voxel FWE-corrected] and there was an interaction with Extinction manipulation (eye, no-eye) [*p*<.003 peak-voxel FWE-SVC]. We found dACC responses for the no-eye movement manipulation [*p*<.007 peak-voxel FWE-corrected], but no differential responses for the eye movement manipulation was present. This is in line with previous findings (Schiller et al., 2013) and could be because this analysis reflects both the process of recovery as well as the process of re-extinction.

## Discussion

This study aimed to test the hypotheses that goal-directed eye movements, as used in Eye Movement Desensitization and Reprocessing (EMDR) therapy, can enhance extinction through amygdala suppression. First, we found that goal-directed eye movements (Experiment 1 and 2) as well as a working memory task (Experiment 1) deactivated the amygdala. Second, we found that both tasks (Experiment 1) altered connectivity between the amygdala and the dorsal fronto-parietal network as well as connectivity between the amygdala and the ventromedial prefrontal cortex. Third, a precisely timed application of the eye movements during extinction learning blocked spontaneous recovery 24h later (Experiment 2). Fourth, while fear responses on average recovered after reinstatement, recovery was attenuated when participants had stronger amygdala deactivations during eye movements (Experiment 2). Given that we found similar amygdala suppression in another task taxing working memory (Experiment 1), the reported effects on fear recovery are likely not specific to eye movements. Furthermore, since fear recovered following reinstatement, our findings provide no evidence that execution of eye movements induces unlearning.

A potential explanation for *why* amygdala deactivation occurs, is that large-scale brain networks act reciprocally (Fox et al., 2005) and compete for resources (Hermans et al., 2014). Acute stress engages the amygdala but impairs dorsal fronto-parietal network functioning (Hermans et al., 2014). Our data confirm that engaging the dorsal fronto-parietal network has the opposite effect of deactivating the amygdala. We furthermore found coupling between the amygdala and dorsolateral as well as ventromedial prefrontal pathways. The vmPFC plays a crucial role in extinction learning and thought to mediate dorsolateral PFC-amygdala interactions (Delgado et al., 2008). This finding aligns closely with the literature on cognitive regulation of emotion (Ochsner and Gross, 2005), while revealing that mere eye movements are sufficient to engage these pathways.

A consequence of this resource competition might be that fear expression is attenuated. Startle responses, for instance, are reduced when performing a working memory task (Vytal et al., 2012) and patients with amygdala lesions show enhanced working memory performance (Morgan et al., 2012). If this mechanism underlies the role of eye movements in reducing traumatic symptoms, then any task taxing working memory should have similar effects. Indeed, emotionality and vividness of autobiographical memories is reduced when memory reactivation is paired with working memory tasks (Holmes et al., 2009; Engelhard et al., 2010; James et al., 2015). Lastly, other types of cognitive control, such as emotion regulation, suppress amygdala activity and alter emotionality during autobiographical memory recollection (Denkova et al., 2015). Our data therefore provide an explanation for how both eye movements and tasks involving cognitive control could affect the emotionality of memories.

Spontaneous recovery was diminished after extinction with eye movements. The dominant view on post-extinction recovery (Maren, 2011; Dunsmoor et al., 2015b) holds that this can be due to updating the original CS-US association or to the formation of a stronger new extinction memory. In line with the latter account, differential fear responses recovered after reinstatement, indicating the CS-US association was not fully eliminated. A similar reduction in spontaneous recovery was observed in a study in which the US was replaced by a non-aversive tone during extinction (Dunsmoor et al., 2015a). One possibility, therefore, is that eye movements following the CS presentation, similar to a tone, strengthen extinction by reducing the ambiguity of the CS either predicting the US or not predicting anything. Unlike a tone, however, eye movements suppress amygdala activity and possibly attenuate fear responses (Vytal et al., 2012). This may allow for additional learned controllability over conditioned responses via subsequent suppression. This interpretation aligns with findings of reduced spontaneous recovery in rats when trained to actively avoid the US during extinction learning (Moscarello and LeDoux, 2013).

The amygdala is, additionally, crucially involved in encoding the CS-US association (Maren, 2011). Amygdala suppression following reactivation could therefore also have led to updating of the CS-US association (*e.g.,* as less aversive) rather than only facilitating new learning. However, we did not find evidence in favor for this explanation. First, we used a design that included multiple presentations instead of a single reminder, necessary to disentangle unlearning from new learning (Eisenberg et al., 2003). Second, fear recovered after reinstatement, indicating full unlearning did not take place. Third, computational modeling analysis revealed eye movements did not prompt a stronger preference for a learning pattern associated with unlearning (Gershman and Niv, 2012; Gershman and Hartley, 2015) We thus propose that eye movements during extinction learning may affect fear recovery by enhancing extinction via newly learned instrumental control over CS-evoked fear responses following memory reactivation (Moscarello and LeDoux, 2013), through pathways engaged during cognitive regulation of emotion (Ochsner and Gross, 2005).

The observed amygdala suppression was located towards the dorsal rather than ventral part. However, we are hesitant in assigning this deactivation to a specific subregion. A comparison between dorsal and ventral is difficult due to inherent problems of gradient EPI sequences, such as signal loss or distortions, which are increased towards the ventral part of the brain (Merboldt et al., 2001; Sladky et al., 2013). These inherent problems (*i.e.*, distance between a brain area and the head coil or cavities) cannot be fully resolved (Merboldt et al., 2001; Sladky et al., 2013). Whether the effect we observed can be attributed to a specific subregion of the amygdala therefore remains an open question.

We found anticipatory FEF activation in response to the CS coupled with eye movements before eye movements took place. This is in line with anticipatory responses observed using electrophysiological recordings in monkeys (*e.g.*, Zhou and Thompson, 2009). In our study, the CS+_eye_ and CS-_eye_ always predicted an eye-movement block, therefore anticipatory responses could be expected. Only five out of 24 participants reported the association between the CS and the eye-movement blocks. Excluding these participants did not affect the results. The FEF activation, moreover, did not predict the reduction in spontaneous recovery, suggesting the anticipatory responses in the FEF did not affect fear recovery.

Despite EMDR being an evidence-based therapy (Bisson et al., 2013; Lee and Cuijpers, 2013), it has received substantial criticism (Devilly, 2002; Rogers and Silver, 2002). Our results shed new light on the working mechanisms of this treatment. One account of EMDR holds that, unlike exposure therapy (Maren, 2011; Bisson et al., 2013), EMDR induced unlearning (Shapiro, 1989; Devilly, 2002; van den Hout and Engelhard, 2012). We found evidence clearly speaking against this claim, since fear recovered following reinstatement. Another controversy regarding EMDR concerns the role of eye movements, which some regard as crucial (Shapiro, 1989), while others argue they have no added value (Rogers and Silver, 2002) or merely serve as a distractor (Devilly, 2002). Our data demonstrate that eye movements have added value above standard extinction learning. However, the data from Experiment 1 suggests that any task taxing working memory would suppress amygdala activity and have similar effects. Indeed, there is a large body of research indicating that working memory tasks reduce the emotionality of memories as well (Holmes et al., 2009; Engelhard et al., 2011; James et al., 2015; Iyadurai et al., 2017). These manipulations have also been shown to be effective in a clinical setting (Iyadurai et al., 2017). Moreover, only tasks with a working memory load reduce the emotionality of memories, while visual distraction does not (Onderdonk and van den Hout, 2016). In sum, our data contradict some of the contentious claims that have been made regarding the underlying mechanism of EMDR.

Several limitations regarding our study need to be mentioned. First, some of our findings are only just statistically significant, and were obtained in a limited sample (n = 24 in both experiments). While amygdala suppression due to eye movements was replicated across experiments, the effect on extinction learning was only tested in Experiment 2 and therefore awaits independent replication. Second, the experimental model of EMDR we developed in this study has inherent limitations because it is impossible to capture every aspect of this therapy (*e.g.*, regarding timing of trauma recall and eye movements, or effects of repeated sessions) in a controlled experiment. Future studies should therefore focus on (1) establishing the reproducibility and generalizability of our findings, (2) investigating the specificity of the observed effects on extinction learning to eye movements, and (3) further illuminating the causal chain from taxing working memory to amygdala suppression and enhanced extinction learning.

In conclusion, our findings show eye movements have added value in safety learning above standard extinction alone. This effect, while likely not specific to eye movements, is associated with amygdala deactivation as a consequence of reciprocally coupled activation of the dorsal fronto-parietal network, likely via ventromedial prefrontal pathways similar to those involved in cognitive regulation of emotion. A key advantage of amygdala deactivation through behavioral manipulations, rather than via pharmacological treatments, is that they are non-invasive, precise in time and duration, and shown to be clinically effective (Bisson et al., 2013). Our findings provide a parsimonious account for how a wide range of behavioral manipulations including working memory tasks, a game of Tetris and eye movements can alter retention of emotional memories.

## Acknowledgments

We thank Sasha Bouman, Frederiek Wijers, and Juul Willering for expert advice on EMDR and Laura van Weerdenburg for help with data collection. This work was supported by grants from the EMDR Research Foundation and the European Research Council (ERC-2015-CoG 682591). Funding sources had no role in study design, data analysis, or writing of the manuscript.

